# Feedforward and feedback population dynamics during binocular conflict in mouse visual cortex

**DOI:** 10.1101/2025.10.02.679998

**Authors:** Melina Timplalexi, William M. Connelly, Adam Ranson

## Abstract

Binocular rivalry arises when incongruent images are presented to the two eyes, producing stochastic alternations in perceptual dominance. While rivalry has been extensively studied in species with highly developed binocular vision, it is unclear whether similar representational dynamics occur in the mouse, a model system that allows large-scale cellular and circuit-level measurements. Here we used two-photon calcium imaging in awake mice to examine the population dynamics of primary visual cortex (V1) and long-range feedback axons from retrosplenial cortex (RSC) during presentation of dichoptically incongruent drifting gratings and natural movies. At the single-cell level, incongruent stimulation increased trial-to-trial variability of visually evoked responses relative to monocular stimulation. Linear SVM decoders trained on monocular responses revealed that during prolonged incongruent stimulation, V1 population activity alternated stochastically between representations of the two competing stimuli in a contrast-dependent manner. Decoder confidence was independent of pupil-indexed arousal state suggesting the dynamics observed may depend mostly on feedforward mechanisms. Transition analyses showed that switches in decoder output were typically driven by the emergence of responses to the ipsilateral stimulus, consistent with release from suppression of the non-dominant population. During incongruent presentation of natural movies, similar representational alternations were observed, indicating that rivalry-like dynamics were not dependent on orientation-selective adaptation. Imaging of RSC→V1 feedback axons revealed retinotopically specific, eye and orientation-selective signals that also alternated in dominance across time. These results establish the mouse as a model of rivalry-like cortical dynamics, demonstrate that both feedforward and feedback circuits contribute to representational alternation during binocular conflict, and provide a framework for mechanistic dissection of bistable perception.

## Introduction

Binocular rivalry occurs when incongruent stimuli are presented to corresponding binocular visual field positions of the two eyes (*1–3*). Instead of perceiving a fusion of the two images, the stimuli compete for perceptual control, resulting in a percept which changes stochastically, with one or other of the stimulus often dominating for several seconds. In human and non-human primates, diverse factors bias the bistable alternation which occurs during rivalry. These include ‘stimulus-intrinsic’ attributes such as contrast and spatio-temporal characteristics, and ‘extrinsic’ factors such as stimulus context or attention of the observer which are thought to be conveyed via cortical horizontal or feedback circuitry.

Neural mechanisms of rivalry have been mostly studied in the visual system of mammals with highly developed binocular vision such as humans, monkeys and cats, and neural correlates of the currently perceived stimulus have been identified at multiple stages of visual processing including from the LGN (*4, 5*), primary visual cortex (*6–11*), MT (*12, 13*), fusiform face area (*14, 15*), and depend on the nature of the stimuli in conflict. However, these species are not amenable to many of the advanced experimental approaches available in mice which would permit larger scale neutral recordings and a more detailed circuit level understanding of cortical computations underlying rivalry. Although mice lack some characteristic organizational features of more developed visual systems such as ocular dominance columns and orientation pinwheels (although see *16–18*), they possess a well characterized binocular visual cortical region (*19–21*), and as in cats and monkeys, retinal inputs from the binocular visual field primarily functionally converge in the primary visual cortex (*22*). Mice also possess increasingly well understood cortical feedback circuitry though which extrinsic factors modulate sensory processing (*23*) – this includes both short-range feedback from higher visual areas (*23–28*) and long-range feedback from frontal regions (*29–33*) and retrosplenial cortex (*23, 34, 35*). Despite these parallels in visual system organisation, it remains largely unknown whether mice exhibit rivalry-like processing when presented with binocularly incompatible stimuli. Studies of processing of binocular incongruent visual stimuli in mice have been limited to examining the encoding of binocular disparities of the same stimulus (*36–38*) or in a recent report during contrast reversing stimuli which provide limited insight into rivalry which exhibits dynamics which play out over time (*39*).

Here we aim to examine the mouse visual system as a model to study temporal dynamics and circuit mechanisms of binocular rivalry using a no-report paradigm in which representations are decoded directly from neural activity rather than via a behavioral response (*40*). We use two-photon calcium imaging in mouse binocular V1 during mismatched short and long presentations of drifting gratings and natural movie stimuli in awake mice, combined with population decoding, to examine temporal dynamics of representations elicited by binocularly incongruent stimuli. To assess the impact of stimulus extrinsic factors on conflict processing, we additionally use pupillography to index arousal, and functional axonal imaging to measure activity of visual feedback signals from retrosplenial cortex under the same conditions. Decoding analysis of V1 activity revealed rivalry-like stochastic trial-to-trail and moment-to-moment alternations of stimulus dominance during incongruent stimulus presentation, which as observed in other species depended strongly on stimulus contrast, but which did not depend on arousal state. Analysis of neural subpopulations encoding the two competing stimuli at moments of transition in decoded dominant stimulus, revealed alternations were mostly driven by brief breakthrough of the usually suppressed responses to the ipsilateral eye. RSC feedback axons were found to relay retinotopically specific visual signals back to binocular V1, which decoding analysis showed were more selective to stimulated eye then stimulus orientation. Together these results provide new insights into population dynamics of both V1 and feedback circuits and establish a novel paradigm for understanding the circuit basis of bistability during stimulus conflict.

## Results

### Binocularly conflicting stimuli elicit unstable V1 representations

We first aimed to determine how briefly presented binocularly mismatched stimuli are represented in the mouse primary visual cortex (V1), and in particular whether alternation between competing neural representations could be observed from trial to trial. Neurons in the binocular segment of V1 were labelled with a genetically encoded calcium indicator using a stereotaxically targeted AAV injection. Dichoptic stimulus presentation in the binocular visual field was performed using a mirror system which allowed stimuli to be presented independently to the two eyes on two separate screens (Figure 1A). Retinotopic mapping was first used to identify layer 2/3 neurons in V1 representing binocular visual space, and to optimize stimulus elevation to match the receptive field of recorded neurons. One of two orthogonally oriented drifting grating stimuli was then presented, either to one eye in isolation or to both eyes. When presented to both eyes, the stimulus orientation either matched or mismatched, producing a congruent or incongruent stimulus.

**Figure 1.**
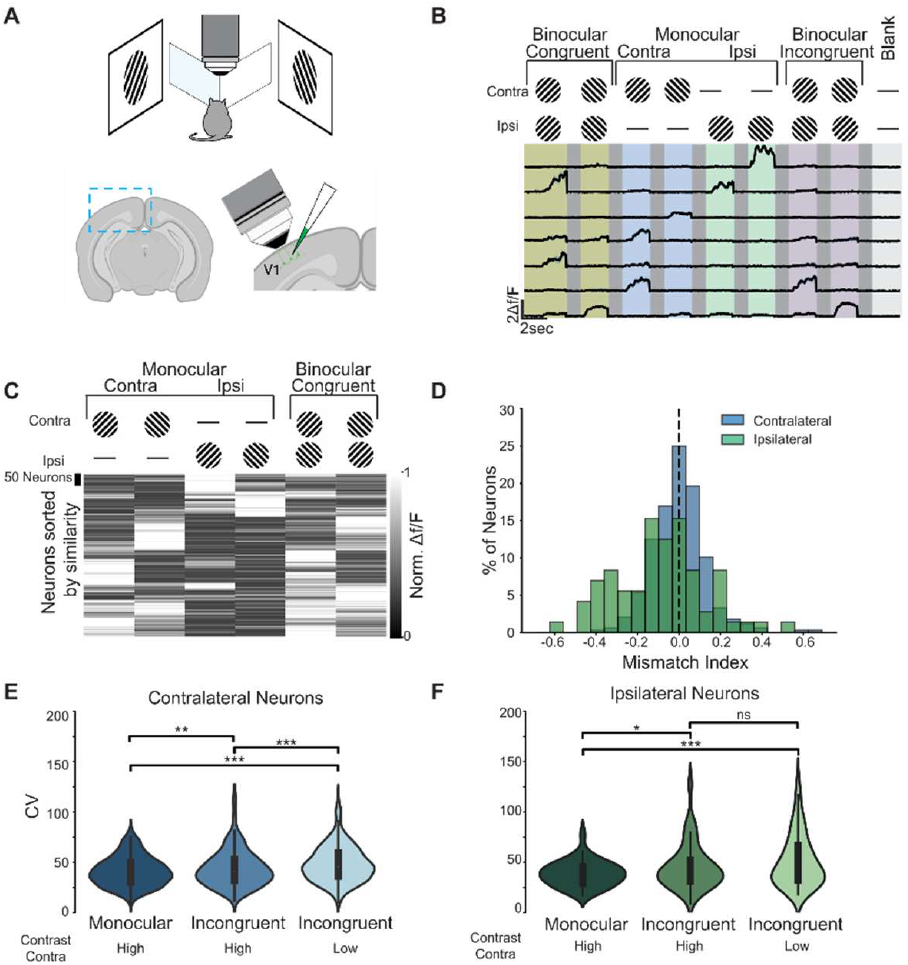
Binocular integration of congruent and incongruent stimuli in single V1 neurons. (A) Schematic representation of the mirror setup for dichoptic stimulation (upper) and injection/recording site (lower). (B) Example neurons recorded under different stimulus conditions. Animals were presented with 45° and 135° oriented stimuli for 2 seconds under monocular, binocular congruent, or binocular incongruent conditions, with an additional blank condition. (C) Normalized neuronal activity in the binocular congruent and monocular conditions for both orientations. (D) Distribution of mismatch indices for contralateral and ipsilateral neurons. (E) Coefficient of variation across all neurons comparing monocular and incongruent conditions at 80% or 40% contrast of the stimulus applied to the contralateral eye. As in E but for ipsilateral neurons. n = 2176 neurons from 7 experiments from 6 mice.

As previously reported in mouse binocular V1, within each field of view within the more lateral segment of V1 which encodes binocular visual space, we found intermingled binocularly and monocularly responsive neurons (Figure 1B; uniquely monocular/contralateral eye responsive: 65.0%; uniquely monocular/ipsilateral eye responsive: 13.9%; binocular: 21.1% of the visually responsive neurons) (*41–43*). For brevity, here we term neurons which respond to stimulation of binocular visual space of only one eye, either ‘contralateral’ or ‘ipsilateral’ neurons, depending upon the eye they respond to, with ‘binocular’ neurons referring to those which respond to stimulation of both eyes. The overall greater occurrence of contralateral as compared to ipsilateral neurons is as previously reported and is consistent with the approximate 2:1 contralateral eye dominance of population responsiveness of neurons in mouse binocular visual cortex and driven in part through the overrepresentation of contralateral eye input to the LGN (*44, 45*).

In general, cells that responded to monocular stimulation of each eye in isolation had an inter-ocularly matching preference for one of the two orthogonally oriented stimuli presented (*46*). To visualize the subtypes of responses of the population to monocular and binocular-matched stimuli we calculated the average response magnitude of each neuron to each stimulus type and sorted the neurons based on response similarity across all stimulus classes. This showed diversity in response types which when clustered by similarity showed four visually identifiable groups of neurons which corresponded to neurons which differed in preferred eye and preferred orientation tuning (Figure 1C).

We next examined responses to incongruent stimuli. To investigate how contralateral and ipsilateral neurons (i.e. neurons which respond to one eye or the other) preferred stimulus responses are modulated on average by congruent or incongruent stimulation of the opposite eye, we calculated a mismatch index in which positive and negative values indicate a preference for matched or mismatched opposite eye stimulation. This measure provides an indication of how a neuron is influenced by being driven by its preferred stimulus in a congruent or incongruent context. Contralateral neurons exhibited a slight positive skew in mismatch index, suggesting that mean activity during congruent trials was higher than during incongruent trials (M=0.00, SD=0.13, skewness=0.72, Figure 1D). In contrast, ipsilateral neurons exhibited a slight negative skew, indicating that mean activity during congruent trials was marginally lower than during incongruent trials (M=−0.08 SD=0.23, skewness=−0.20, independent samples t-test, t = −2.02, p = 0.044, Figure 1D). This suggests ipsilateral neurons are more suppressed during mismatched stimulation of the opposite eye than contralateral neurons.

We reasoned that if a rivalry-like process was occurring at the level of V1 during incongruent stimulation, then response amplitudes of individual neurons would be expected to vary more from trial to trial during incongruent stimulation than during congruent stimulation (i.e., as one stimulus representation or the other dominates). To test this, we calculated the coefficient of variation (CV) of responses of each visually responsive neuron in each trial type. An increase in CV indicates more trial-to-trial variation of a neuron to a given stimulus, while accounting for variation in overall response amplitudes across conditions.

For contralateral neurons, a mixed-effects model revealed a significant main effect of condition, such that CV was higher during both incongruent conditions compared to monocular stimulation. Specifically, compared to monocular stimulation, CV was significantly higher during the incongruent condition of high contralateral contrast (z = 2.63, p = .009,) and the incongruent condition of low contralateral contrast (z = 6.79, p < 0.001, Figure 1E). For ipsilateral neurons, a similar effect was observed. Relative to monocular stimulation, CV was significantly greater during the incongruent condition of high contralateral contrast (z = 2.19, p = 0.028,) and the incongruent condition of low contralateral contrast (z = 3.78, p < .001, Figure 1F). Together, these results demonstrate that stimulus incongruency increases single-cell response variability in V1, and this effect is especially pronounced when the contralateral input is disadvantaged by reduced contrast.

### Dominant population-level representation alternates stochastically during prolonged incongruent stimulation

Given the single-cell short timescale observations above, we next aimed to determine what is globally encoded during incongruent stimulation at the V1 population level. Specifically, we aimed to test if population activity observed during incongruent stimulation alternates (in a rivalry-like manner) between the two activation patterns produced by monocular stimulation with the individual orthogonally oriented constituent stimuli. Previous studies in humans and non-human primates have found that binocular rivalry occurs over extended periods of time, with one of the competing visual representations often dominating perception for several seconds (*6, 14, 15, 47*). In contrast, during short stimulation fusion of representations often occurs (*48*). We next therefore investigated the temporal dynamics of stimulus representations over prolonged periods of continuous stimulation. On each trial we presented oriented drifting gratings to one or both eyes for 30 seconds, which were either of a single orientation (congruent), or of mismatched orthogonal orientations (incongruent).

To decode the temporal dynamics of the dominant representation from the V1 population during stimulation, a linear SVM classifier was trained using responses to a brief (2 second) monocular stimulus presentation (over which activity was averaged) under the assumption that this would capture the subpopulation of neurons which respond to that specific stimulus. As expected for orthogonally oriented grating stimuli, the classifier had a high level of cross-validated accuracy in decoding grating orientation from trials in which gratings were briefly presented to one eye (average accuracy across all experiments 92.5 ± 2%, Figure 2A). The trained SVM was then used to classify 1 second bins of neural activity during the 30 second monocular or incongruent stimulation trials. We hypothesised that rivalry-like alternation of visual representation during incongruent stimulation would be indicated by alternation of classification combined with an above chance level of decoding ‘confidence’ in classification. To gain a measure of classification confidence we transformed SVM decision values into calibrated probabilities using Platt scaling (*49*).

**Figure 2.**
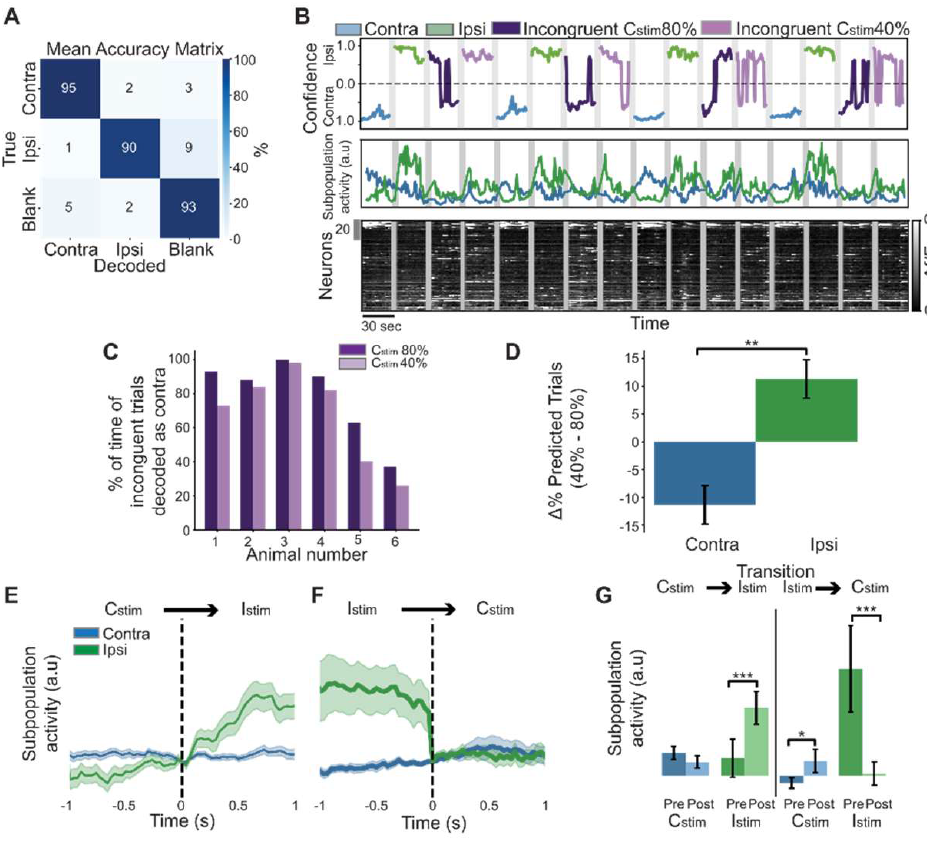
Neuronal population representation during prolonged presentation of incongruent stimuli. (A) Mean confusion matrix of SVM classification of neural activity occurring during monocular trials as having been driven by either a stimulus presented to ipsilateral or contralateral eye or a blank stimulus. (B) Example trials from one mouse showing classifier decision (upper), activity level of contralateral (blue) and ipsilateral (green) encoding subpopulations (middle; derived from SVM classifier weights, see methods), and raster map of activity of all neurons (bottom). (C) Fraction of bins within incongruent trials which were decoded as contralateral or ipsilateral stimulus in each mouse, and how this is affected by reducing contralateral stimulus contrast. (D) Summary of effect of reducing contralateral stimulus contrast on fraction of bins within incongruent trials which were decoded as contralateral or ipsilateral stimulus. (E) Activity level of contralateral and ipsilateral encoding subpopulations aligned to classification decision switches from contralateral to ipsilateral. (F) As in E but for switches from ipsilateral to contralateral. (G) Average pre and post-switch activity levels of contralateral and ipsilateral encoding subpopulations. n = 1977 neurons from 6 experiments from 5 mice.

Since we aimed to use the SVM decoding approach to assess dominant representations over the 30 second trial period, we first sought to validate the stability of decoding over this timescale. We first assessed the dynamics of monocular (i.e. non-conflicting) stimulus presentation during prolonged stimulation to determine if stimulus identity could be stably decoded across all epochs of single trials. While the stimulus class was almost always correctly decoded (monocular average accuracy = 98.3 ± 4.1%), decoding confidence did fluctuate substantially within trial (Figure 2B). To quantify this, we compared classifier confidence on the first and last 10 seconds on each 30 second stimulus presentation. This showed that classifier confidence did significantly decrease during contralateral stimulation (from 0.89 ± 0.01 to 0.81 ± 0.04, paired t-test, p = 0.016), while the decrease during ipsilateral stimulation did not reach statistical significance (from 0.88 ± 0.02 to 0.80 ± 0.05, p = 0.068), although classification confidence remained overall well above chance.

We next used this trained SVM classifier to decode the dominant representation in incongruent trials. We found that in each experiment the stimulus decoded by the classifier was generally dominated by either the contralaterally or ipsilaterally presented stimulus (in the majority of cases the former), however many periods could be observed in which the non-dominant representation emerged as dominant (Figure 2B and C). While confidence levels were overall lower in classifying incongruent versus monocular trials, classifier confidence was significantly higher than expected by chance, as assessed by calculating confidence levels in incongruent trials in which cell identities were shuffled (Figure S2B).

Above we describe that trial-to-trial response variability (i.e. CV) is elevated in both ipsilateral and contralateral stimulus responsive neurons when the contralateral population is disadvantaged by reduced contralateral stimulus contrast (Low-contrast contralateral condition). We next assessed whether the total time that the contralateral representation dominates is similarly modulated by relative stimulus contrast. Consistent with this, reducing contralateral stimulus contrast while keeping ipsilateral stimulus contrast fixed led to a decrease in the total duration of incongruent trials decoded as contralateral (Contra: t = –3.51, p = .0085), accompanied by a corresponding increase in the duration decoded as ipsilateral. These results suggest global alternation of dominant representation at the V1 population level during incongruent stimulus presentation which is modulated by stimulus contrast.

### Activity level of contralateral and ipsilateral stimulus encoding subpopulations both drive decoder behavior

We next asked what neural activity dynamics underlie alterations in SVM classification decisions observed during incongruent stimulus presentation. Specifically, we aimed to determine if switches in dominant encoded stimulus (as decoded by the SVM) are characterised by a ramping up of the soon to be dominant population or a ramping down of the soon to be non-dominant population, or whether both processes drive switches in classification. To distinguish between these possibilities, we utilised the feature weights fitted by the SVM, in order to calculate population activity vectors of neurons that are informative of the presence of the contralateral or ipsilaterally presented stimuli, resulting in C_stim_ and I_stim_ neural subpopulation activity vectors respectively (Figure 2B, middle panel). We then aligned these activity vectors to moments at which the classifier decision alternated, either between C_stim_→I_stim_ or I_stim_→C_stim_. We then averaged these weighted activity vectors over the 1 second before and after classification switches. As expected, C_stim_ activity was greater than I_stim_ activity after I_stim_→C_stim_ transitions (Figure 2E), whereas the opposite was the case after C_stim_→I_stim_ transitions (Figure 2F). In the case of C_stim_→I_stim_ transitions, classification change was on average driven by an increase in the I_stim_ encoding population (W = 6, *p* < 0.001), with no significant alteration of the C_stim_ population (W = 231, *p* = 0.984; Figure 2E and G). In contrast, I_stim_→C_stim_ transitions were associated with both an increase in the C_stim_ population (W = 85.0, p = 0.037) and a decrease of the I_stim_ population (W = 28.0, p < 0.001; Figure 2F and G). These data suggest that while both C_stim_ and I_stim_ subpopulations are driving the decision of the SVM decoder, some asymmetry exists in the dynamics underlying C_stim_→I_stim_ vs. I_stim_→C_stim_ transitions. Overall, the dynamics observed are consistent with a gradual easing of suppression of the currently non-dominant population leading up to a switch.

### Natural stimuli also drive stochastic representational switching when presented incongruently

We next aimed to test whether the alternating dynamics we observed during prolonged presentation of incongruent grating stimuli are caused by strongly driving a subpopulation of neurons selective to the orientation in question. We reasoned that grating stimuli may produce adaptation in this subpopulation and result in dynamics specific to synthetic stimuli with a single orientation, spatial, and temporal frequency (*50*). To investigate this possibility, we instead presented mice with natural video stimuli (Figure 3A). In this condition, the incongruent stimulus consisted of the same movie presented to the two eyes, but out of temporal phase by 10 seconds (movie_Δ0s_ and movie_Δ10s_). Neural activity from monocular presentation of the movie to either the contralateral (movie_Δ0s_) or ipsilateral (movie_Δ10s_) eye was used to train a support vector machine (SVM) to classify neural activity as having resulted from movie_Δ0s_ or movie_Δ10s_ (indicating contralateral or ipsilateral stimulus respectively). Neural activity was averaged over 3-second bins for analysis. The classifier achieved an average 86% ± 4% accuracy in predicting whether movie_Δ0s_ or movie_Δ10s_ was occurring, across ten 3-second bins (Figure 3B, ‘Data’). As expected, a control analysis, in which trial identities were randomly shuffled 1000 times, yielded an average accuracy of 48% (Figure 3B, ‘Shuffled’), consistent with chance performance at binary classification. Comparison of movie_Δ0s_ vs. movie_Δ10s_ decoding accuracy over the ten 3 second epochs indicated that neural activity associated with each epoch was not distinguished by the classifier with differing levels of accuracy (repeated-measures ANOVA, main effect of epoch: F(9,45) = 1.78, p = 0.10). We next used the monocularly trained classifier to decode neural activity in incongruent trials and found that similar to grating stimuli, the decoded stimulus often alternated between the typically dominant contralateral stimulus and the ipsilateral stimulus (Figure 3C). The average fraction of each incongruent trials classified as dominant eye vs. non-dominant eye was comparable between grating and natural stimuli suggesting that grating driven adaptation is not a critical driver of alternation.

**Figure 3.**
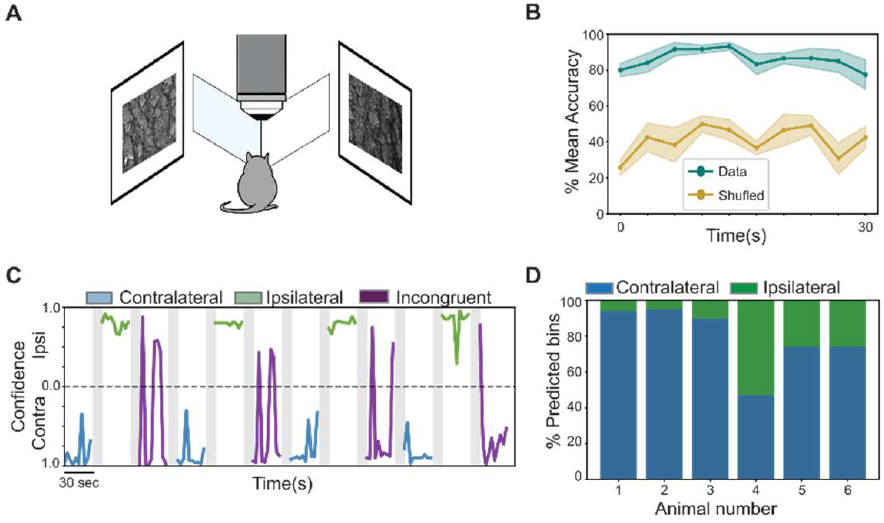
V1 population dynamics during natural video stimulus. (A) Schematic of visual stimulation. (B) Mean SVM decoding accuracy (decoding whether movie_Δ0s_ is being presented to ipsilateral eye or movie_Δ10s_ is being presented to contralateral eye) based on neural activity (actual data, green, or shuffled, orange) in each of ten 3 second bins. (C) Example trials from one mouse showing decoding of stimulus type from 1 second bins of neural activity for each stimulus type. (D) Fraction of 3 second neural activity bins decoded as contralateral or ipsilateral stimulus in each mouse during movie stimulus. n = 845 neurons from 6 experiments from 6 mice.

### Arousal state does not modulate incongruent stimulus processing in V1

We next considered whether processing of incongruent stimuli may also depend on the extrinsic factor of the animal’s state of attentiveness. To evaluate this, in a subset of experiments we measured pupil diameter as a proxy of arousal state (*51–53*). We found pupil diameter varied over long and short timescales over the course of each 45-minute recording (Figure 4A and B). We first asked if stimulus incongruence itself could drive changes in arousal, since the binocularly mismatched stimulus itself may be surprising to the animal. To quantify stimulus driven pupil diameter changes, we plotted pupil diameter aligned to either monocular or incongruent trial onset and normalised to the pre-trial period and found no evidence of stimulus onset driven pupillary response in either case (quantified as average of 2 second period after stimulus onset; monocular: *M* = –0.255 a.u., *t* = –1.80, *p* = .076; incongruent: *M* = –0.078 a.u., *t*= –0.45, *p* = 0.655; both compared against zero; Figure 4C and D).

**Figure 4.**
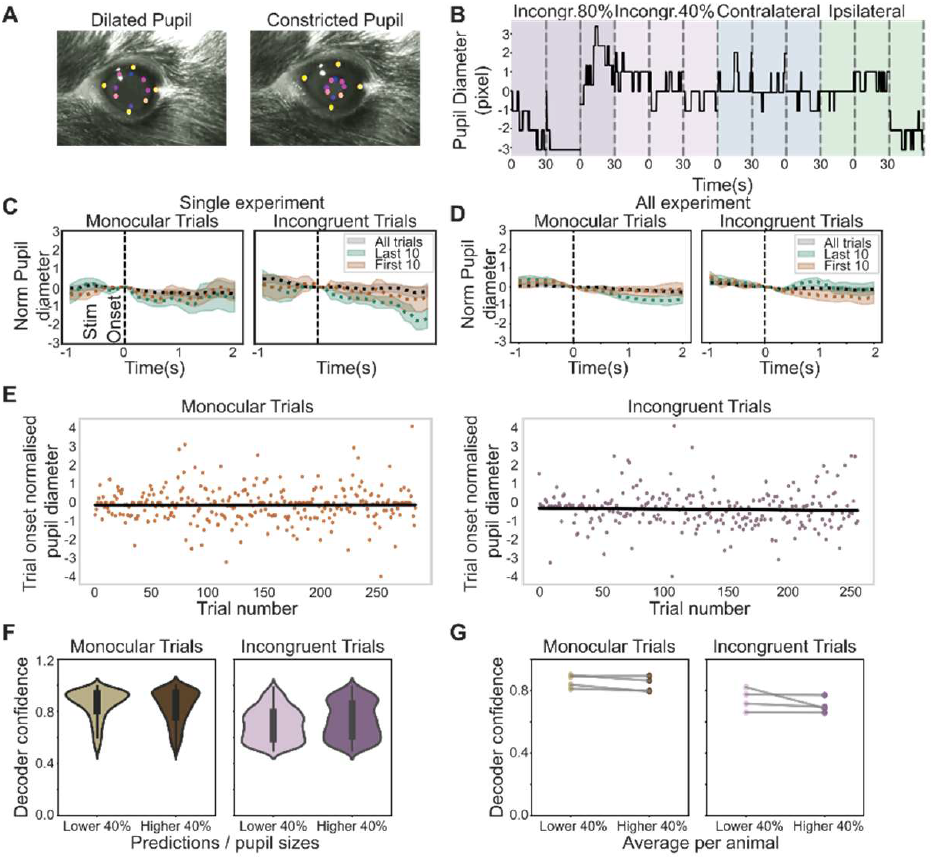
Pupil dilation is insensitive to incongruent stimulus presentation and does not predict representation dynamics. (A) Detected pupil features. (B) Pupil dynamics in an example experiment during different trial types. (C-D) Averaged trial onset aligned and normalised pupil diameter, for one experiment (C) or across all experiments (D). (E) Pre vs. post trial onset pupil change magnitude as a function of trial number on monocular (upper) or incongruent (lower) trials. (F) SVM decoder confidence on monocular (left) and incongruent (right) trials in which the pupil diameter is in the upper or lower 40^th^ percentile. (G) As in F but showing average decoder confidence across all trials of each mouse. n = 1977 neurons from 6 experiments from 5 mice.

Since stimulus driven changes in arousal may diminish with repeated incongruent stimulus presentations (i.e. due to reduced surprise), we additionally compared the pupil response at onset of the first 5 monocular and incongruent trials to the average of the last 10 and again observed no systematic difference (Figure 4C, single experiment, and average across experiments in D). Similarly, analysis of pupil response in all animals as a function of trial number showed no systematic relationship between stimulus onset driven pupil dilation and trial number for either monocular or incongruent stimuli (Figure 4E). Together these data indicate that in this context, stimulus onset per se (either monocular or incongruent) does not drive a pupil response.

We next asked if incongruent stimuli are processed distinctly in states of high vs low arousal, and specifically whether states of high classifier confidence (in which representations more closely resemble one or other of the incongruently presented stimuli and are thus more disambiguated) occur more frequently in high or low arousal states. To test this, we compared classifier confidence in trials in which the average pupil diameter was in the lower vs higher 40th percentile and observed no significant systematic difference (Figure 4F and G). Together these results indicate that arousal state, as indexed by pupil diameter, is neither significantly driven by the incongruent stimuli used in this study, nor is it associated with alterations in how incongruent are disambiguated, suggesting a passive mechanism independent of current processing demands.

### RSC feedback axons relay retinotopically targeted eye specific signals during incongruent stimulation

Top-down contextual information and attentional signals relayed by cortico-cortical feedback are a further proposed modulator of representational alternation during rivalry (*2, 3*). We hypothesised that as conflicting visual signals propagate through visual areas, representations could progressively become disambiguated (and more closely match to subjective perception *54*) as they undergo competition at each stage of visual processing, and that such signals could be fed back to V1 to bias conflicting stimulus resolution. To examine this possibility we labelled neurons in the retrosplenial cortex (Figure 5A), a medial association area that is a major source of long-range sensory feedback to sensory cortex (*23, 34, 35*), and then measured signals fed back to V1 by imaging RSC→V1 feedback axon activity in layer 1 of V1 (Figure 5B). Consistent with recent findings (*35*), we found that a large fraction of RSC→V1 axons were visually responsive and retinotopically tuned, and included a subpopulation relaying feedback signals to V1 neurons specifically encoding the binocular visual field (Figure 5C).

**Figure 5.**
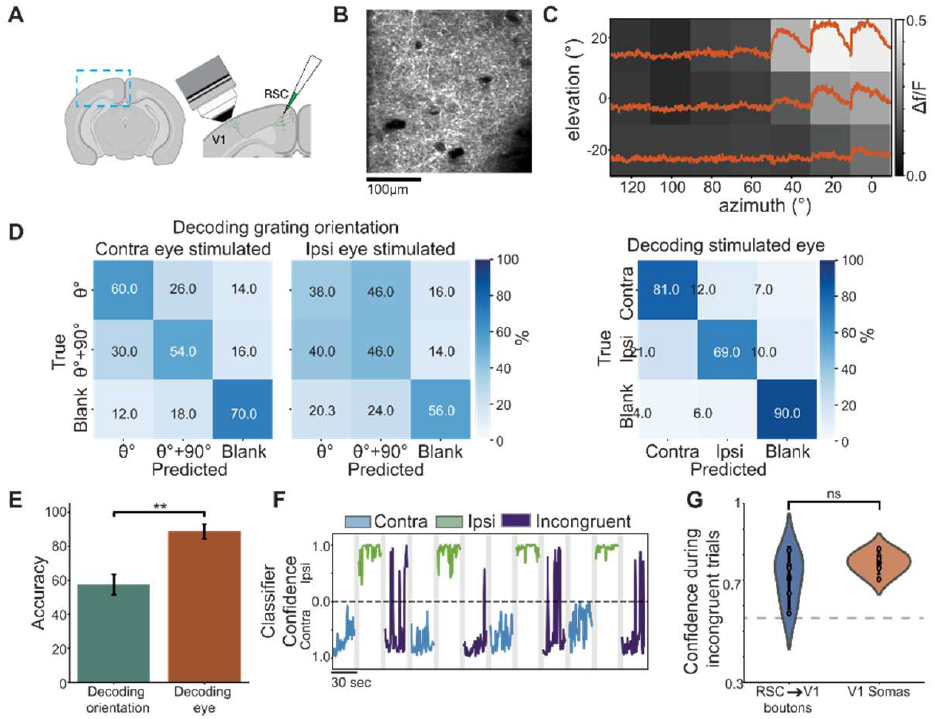
Feedback signals from retrosplenial cortex to V1 are eye specific and retinotopically targeted. (A) Schematic showing labelling of RSC→V1 axons by injection of AAV into RSC. (B) Labelled RSC→V1 axons visualised *in vivo* in layer 1 of V1. (C) Example spatial tuning of a binocular visual space encoding RSC→V1 bouton. (D) Mean confusion matrices from SVM classification of stimulus orientation from RSC→V1 bouton activity occurring during monocular stimulation of contralateral eye (left) or ipsilateral eye (middle), or classification of stimulated eye (right). (E) Average SVM classification when decoding stimulus orientation or stimulated eye from RSC→V1 bouton activity. (F) Example trials from one mouse showing decoding of stimulus type from 1 second bins of neural activity for each stimulus type. (G) Comparison of SVM decoding confidence when decoding from RSC→V1 boutons vs. V1 somas. n = 615 boutons from 5 experiments from 4 mice.

RSC→V1 feedback is largely untuned at the single bouton level to grating stimulus orientation during passive viewing (*34, 35*), despite reports that the RSC soma population more widely exhibits some orientation selectivity (*55, 56*). If RSC→V1 feedback were to modulate V1 responses during interocular conflict, a prerequisite would be some degree of orientation or eye selectivity of RSC→V1 bouton responses at the population level. We therefore next asked if grating stimulus orientation or stimulated eye could be decoded if the overall measured RSC→V1 feedback population was used for decoding. As in the analysis of V1 responses, we trained an SVM to decode either stimulus orientation or stimulated eye from neural activity from all RSC→V1 boutons detected in V1. Classifiers achieved well above chance decoding of stimulus orientation during contralateral eye stimulation (61% average cross validated performance, while chance level is 33%, Figure 5D left), while decoding during ipsilateral eye stimulation was lower (46.66% average cross validated performance, Figure 5D middle). Unexpectedly, we also found the classifier was able to more accurately decode stimulated eye from RSC→V1 bouton activity than orientation tuning (80% average cross validated performance, t = −6.063 p = 0.009; Figure 5D, right and 5E), indicating eye representations are segregated in at least an RSC subpopulation.

Applying this decoder to RSC→V1 bouton activity for 30 second periods of stimulation revealed alternation of the dominant representation during incongruent stimulation (Figure 5F) similar to that observed in V1 soma activity. We tested how classifier confidence in decoding the currently dominant RSC→V1 bouton representation during incongruent stimulation compared to that when decoding V1 representations under identical conditions and found it did not differ significantly (t = 1.235, p = 0.2692; Figure 5G). This finding is not supportive of progressive disambiguation over the rodent dorsal visual processing stream, at least in the subset of RSC neurons which provide feedback to V1.

These results show that RSC→V1 axons feedback retinotopic eye specific signals to V1, and during incongruent stimulation exhibit alternation of dominance. Previous findings indicate that binocular visual field responsive RSC→V1 boutons selectively target binocular regions of the V1 retinotopic map and could therefore feasibly be a driver of transitions in lower sensory encoding and subsequent perception.

## Discussion

In this study we combined two-photon imaging of V1 population activity and associational cortex feedback from RSC to V1, with dichoptic visual stimulation to probe how conflicting visual stimuli are represented in the mouse visual system. We found that incongruent visual stimulation results in greater trial-to-trial instability of visual evoked activity in single cells, consistent with a rivalry-like process. Supportive of this interpretation, decoding analysis of V1 cellular activity showed that at the population level this translates to coherent alternations of activity patterns between the two presented stimuli. This suggests that the mouse is a feasible model in which to examine aspects of the circuit basis of binocular rivalry.

We observed that the stimulus presented to the contralateral eye almost always dominates the V1 binocular cortical representation. However, as in other species, stimulus ‘strength’ (in this case contrast) could be used to modulate the amount of time one or other of the stimuli dominated. Switches of the currently dominant stimulus (decoded at the population level) were primarily driven by an apparent ramping up or suppression of the weak representation (in this case that driven by the ipsilateral eye). The circuit mechanisms through which this interplay occurs likely involve both locally driven feedforward inhibitory circuits, and inhibition recruited through longer range feedback which targets VIP and SOM interneurons. The mouse presents a model in which these mechanistic questions can be feasibly addressed.

While cortical representations were typically strongly contralaterally dominated, there were fields of view in which this was not the case. This may be due to sampling from neurons within ocular dominance domains which have recently been reported in mice which contain a greater or lesser abundance of ipsilateral eye responsive neurons (*16*). It will be of interest to employ mesoscopic two-photon imaging of V1 during incongruent stimulation, both to confirm that this is the origin of the variation in ocular dominance we see and to test for the existence of competition at a larger spatial scale between these domains as has been reported in other species.

The absence of a relationship in our experiments between arousal state, indexed by pupil diameter, and incongruent stimulus processing, suggests that the dynamics observed may be best described as being determined by a simple feedforward process. Both a strength and weakness of our study is that we use a no-report paradigm that depends on direct representational decoding from neural activity rather than a behavioral report or involuntary optokinetic nystagmus. While this avoids confounds associated with behavioral/motor responses, it is possible that processing of stimulus conflict occurs differently when animals are engaged in active visual discrimination. In the context of a no-report paradigm, it will be of interest to probe visual representations at other more advanced stages of visual processing after V1 to test for evidence of increasingly clear response bistability.

Finally, we provide the first report of feedback from a higher cortical associational area to primary visual cortex during binocular stimulus conflict. Unexpectedly, while feedback activity from RSC to V1 was weakly stimulus orientation tuned, it was more highly eye and retinotopic location specific. Decoding analysis during incongruent stimulation showed that overall decoder confidence was not higher when decoding dominant representation from RSC→V1 bouton activity compared to from V1 somas, inconsistent with representations becoming more bistable over successive stages of cortical processing. However, this metric is likely impacted by many other factors, not least by differences in signal to noise when imaging boutons compared to somas. One recent report suggested interocularly mismatched contrast reversing gratings resulted in reduced activity of SOM interneurons which may increase the influence of feedback to V1. Further experiments with concurrent RSC axon and V1 neuron recordings will be required to determine if RSC axons selective to one eye or the other, target V1 domains which are also dominated by that same eye, and what the impact of this feedback is on V1 activity.

This study presents evidence of a rivalry-like process at a neural level, even at early stages of the mouse sensory system. The results provide a basis on which to apply the powerful experimental approaches only available in rodent models to gain new insights into the circuit elements and connectivity organisation that underlie bistable perception.

## Experimental model and subject details

### Animals

All experimental procedures were carried out under the Ethics Committee on Animal and Human Experimentation from the Universitat Autònoma de Barcelona and followed the European Communities Council Directive 2010/63/EU. Experiments were performed on adult (age>P90) C57BL/6J mice. Experiments were carried out in mice of either sex, group-housed, with 12:12h light-dark conditions, *ad libitum* access to food and water. All recordings were made during the light period.

## Method details

### Animal Surgical Preparation and virus injection

All surgical procedures were conducted under aseptic conditions. Before the cranial window surgery, animals were administered the anti-inflammatory drugs Rimadyl (5 mg/kg, s.c.) and dexamethasone (0.15 mg/kg, i.m.), and antibiotic Baytril (5 mg/kg, s.c.). Anesthesia was delivered in gas form with a 5% concentration of isoflurane in oxygen and then maintained at 1.5-2% for the duration of the surgery. Once anesthetized, mice were placed onto a heating pad to maintain their body temperature at 37°C, and head fixed using ear bars into a stereotaxic frame. The area of the surgery was shaved and cleaned using a 70% ethanol solution followed by an antiseptic iodine solution, and two injections of lidocaine (2% w/v) were made subcutaneously into the scalp for local anaesthesia. The scalp and the periosteum overlying the area of headplate fixation were then resected. A custom head plate was then attached to the skull using Vetbond and secured with dental cement (Paladur, Heraeus Kulzer), with the aperture centered over the binocular area of the right hemisphere V1 (−3.2 posterior and 3.1 lateral from bregma). A 3 mm circular craniotomy was next performed, centered on the stereotaxically identified binocular area of V1.

For experiments in which V1 neurons were labelled, after the V1 craniotomy was performed, an AAV was injected at multiple sites at a depth of 200-400µm to drive expression of jGCaMP7f (pGP-AAV-syn-jGCaMP7f-WPRE, Addgene plasmid #104488, 5 × 10^12^ GC per millilitre titre, volume 100 nL).

For retrosplenial cortex (RSC) injection, a small craniotomy was made above the retrosplenial cortex (−2.8mm posterior from bregma, 0.3mm lateral from the midline) using a dental drill, thinning of the bone and then removal of a small bone flap using a hypodermic needle. After the brain was exposed, an AAV was injected at a depth of 400µm to drive expression of jGCaMP8s (pAAV-hSynapsin1-axon-jGCaMP8s-P2A-mRuby3, Addgene plasmid #172921, 5 × 10^12^ GC per millilitre titre, volume 100 nL). The craniotomy of the RSC was sealed with Vetbond.

Injections were made using either a Nanoject II system (Drummond Scientific Company) or a Stereodrive Drill & Microinjection Robot (Neurostar) at a rate of 10 nL/min using pulled and beveled oil-filled glass micropipettes with a tip outer diameter of approximately 30-40µm. After injections, the craniotomy over V1 was closed with a glass insert constructed from 3 layers of circular glass 100um thickness (1 of 4 mm and 2 of 3 mm diameter) bonded together with UV cured optical adhesive (Norland Products; catalog no. 7106). Habituation to head fixation and imaging started a minimum of 2 weeks after surgery.

### *In vivo* imaging

*In vivo* two-photon imaging was performed using a Movable Objective Microscope with Resonant Scan box (Sutter Instrument) with a N16XLWD-PF objective (Nikon, 0.8 NA, 3.0 mm WD). Imaging of the green calcium indicators jGCaMP7 and jGCaMP8 was performed by excitation at 920nm using either an 80 MHz Ti-Sapphire laser (Chameleon Ultra II, Coherent) or a 920 nm 80 MHz fiber laser (FemtoFiber ultra 920, TOPTICA). Imaging of jRGECO1a was performed by excitation at 1050nm using an 80 MHz fiber laser (FemtoFiber ultra 1050nm, TOPTICA). In all experiments laser power at the sample was limited to approximately 60mW. Data were acquired a frame rate of 30 Hz (512 × 512 pixels) using Scanimage 5 with a standalone NI FLEX RIO DAQ (National Instruments). The field of view size in experiments in which V1 somas were recorded was approximately 250 x 250 µm^2^ while in experiments in which boutons were recorded the field of view was approximately 100 x 100 µm^2^. Synchronisation of imaging with visual stimulation and other data (behavioral and eye cameras, rotary encoder of linear treadmill) was attained by acquiring timing signals on a separate data acquisition card (NI PCIe-6323, National Instruments) using custom MATLAB code (https://github.com/ransona/Timeline-UAB).

### Behavioral and pupil position monitoring

In all the experiments videos of both eyes of the animals were recorded using USB monochrome cameras (Imaging Source model DMK 22AUC03 with lens Azure-7524 mm and a 800nm short pass filter) under IR illumination, with frames acquired using custom Python code (https://github.com/ransona/py_eye) and synchronised using a digital signal produced by an Arduino Uno (Arduino). Eye videos were processed to obtain pupil position by first detecting a number of eye landmarks using DeepLabCut (*57*). Namely, four points were detected which corresponded to the centre of upper and lower eye lid and medial and lateral eye corners, and eight points were detected which corresponded to the edges of the pupil (i.e. an octagon). The model was trained on 600 hand labelled frames and was validated by manual inspection of experiments with detected features overlaid on eye video frames. While the network’s placement was visually indistinguishable from human placement for the vast majority of frames, the network would always attempt to place all eight pupil markers, even if the mouse was blinking or the pupil was otherwise mostly obscured by the eyelids. To remedy this, a program was written to construct an eye shape by passing a pair of parabolic curves through the medial, superior, and lateral and medial, inferior, and lateral eye markers respectively, and any pupil markers that fell outside this eye shape were discarded. In addition, if a set of basic assumptions about the eye shape were violated – for instance, if the medial eye marker was lateral to the lateral eye marker – all pupil markers were discarded. Finally, if six or more pupil markers were valid, an ellipse was fitted to them to minimize the least-squared error. Together these measures resulted in high fidelity tracking of pupil behavior across frames (*29*). Timepoints at which the pupil could not be detected were excluded from further analysis. Animal locomotion on the single axis treadmill was measured using a rotary encoder with 1024 steps per rotation (Kübler, 05.2400.1122.0100) acquired using an NI PCIe-6323 DAQ (National Instruments).

### Visual stimuli

Visual stimuli were generated using a custom written interactive visual stimulation rig (https://github.com/neurogears/ranson-rig) controllable by OSC and built upon Bonsai and BonVision (*58*). Stimuli were presented on three 54 x 40 cm^2^ LED Monitors at 60 Hz refresh rate (D24-20, Lenovo) positioned such that stimuli could be presented from approximately −120° to +120° in azimuth, and −30° to +30° in elevation. Stimuli were warped to correct for screen flatness using Bonvision and screen luminance was linearised. To avoid contamination of imaging data by stimulus screen light, LED screens were modified in house to flicker the screen LED in synchrony with microscope line scanning such that LEDs were only illuminated during scan line turn around. A custom designed circuit was used to allow control over screen LED on time and maintenance of screen brightness during frame flyback (https://github.com/ransona/screen-LED-blanking-circuit). Stimulus timing was synchronized to other data acquisition using both digital signals produced by Bonsai via a Harp Behavior board (OEPS-1216, open ephys) and a photodiode (OEPS-3011, open ephys) which acquired an on-screen black/white alternating sync square.

Binocular visual space encoding neurons were first localised by performing retinotopic mapping on candidate regions which were expected to be suitable based on stereotaxic injection coordinates with regions encoding approximately the central 40 degrees of visual space during ipsilateral eye stimulation treated as binocular. Retinotopic mapping stimuli consisted of 30 x 30° circular horizontally oriented square wave gratings of spatial frequency 0.05 cycles per degree which drifted at a temporal frequency of 2 Hz. Stimuli were presented in positions on a 7 x 3 grid with stimuli centred in positions ranging from 120° to 0° in azimuth and −20° to 20° in elevation. Each grating stimulus was presented for 2 seconds with a 1 second interstimulus interval and each position was repeated 10 times in a pseudorandom order.

Having identified a region as encoding binocular visual space, silver coated mirrors at right angles to each other were positioned directly in front of the mouse, as close to the head as possible without direct contact (see Figure 1A). Barriers were placed directly lateral to the eyes to prevent direct viewing of stimuli on laterally placed screens. The same retinotopic mapping stimulus described above was then used to confirm the binocular region was now activated by stimulation on the area of the screen to the left and right of the mouse. Left and right stimulus position was optimised independently to maximise responses of binocular visual space encoding neurons.

In experiments in which conflicting or matched oriented gratings were presented (Figure 1, 2, 4 and 5), stimuli consisted of 30 x 30° oriented (either 45° and 135°) circular square wave gratings of spatial frequency 0.05 cycles per degree which drifted at a temporal frequency of 2 Hz, as well as a blank stimulus. Each grating stimulus was presented for 2 seconds (used for characterisation of basic response properties in Figure 1, and for SVM decoder training) or 30 seconds (used for decoding during congruent or incongruent conditions) with a 1 second interstimulus interval and each stimulus was repeated 8 times in a pseudorandom order. In experiments in which natural video was presented (Figure 3), movie stimulus consisted of a 40 second drifting natural bark texture, the first 30 seconds of which was presented to one eye (movie_Δ0s_) and the last 30 seconds of which was presented to the other eye (movie_Δ10s_).

### Calcium imaging data processing

Calcium imaging data was registered and segregated into neuronal regions of interest (i.e. RSC boutons or V1 somas) using Suite2P (*59*). The time series of each ROI was then converted from a raw fluorescence value to ΔF/F with the denominator F value trace constructed by calculating the 5th percentile of the smoothed F value within a 20 second window centred on each sample in the F trace.

## Quantification and statistical analysis

### Analysis of visual response selectivity of single V1 neurons

In basic visual response characterisation experiments (Figure 1), neurons were classified as visually responsive on the basis of an ANOVA conducted across responses to all stimulus locations and the blank, with a significance threshold of p < 0.05 applied to determine responsiveness, as well as requiring that a bouton has a mean response to preferred stimulus which was greater than 2 standard deviations above the average blank response. Average response amplitude to stimulation in each position was then calculated for each visually responsive neuron/bouton with response calculated by averaging ΔF/F between 0.5 and 2 seconds after stimulus onset. In order to sort neurons by response similarity (Figure 1C) we applied hierarchical clustering (Ward’s method) to the pairwise distances and reordered the matrix according to the optimal leaf order. The mismatch index (Figure 1D) was calculated using the following equation:

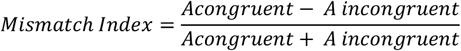

where A denotes the mean neuronal activity in each condition. Here, positive and negative values indicate a preference for matched or mismatched opposite eye stimulation.

To assess whether the population response to incongruent stimuli was simply the sum of the responses to the component monocular stimuli, we computed the squared Mahalanobis distance 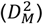 between the summed mean responses and the incongruent response. Under the null hypothesis that the responses during incongruent stimuli is equal to the sum of the responses to the component stimuli, this distance zero and 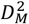 follows a chi-squared distribution with degrees of freedom equal to the number of neurons. The activity was normalized using a Yeo-Johnson transform.

In order the limit this analysis to neurons directly related to the task, an L1-regularized linear SVM was fitted to the neural responses to classify stimuli as either contralateral or ipsilateral. The regularization was applied as weakly as possible, while still maximizing accuracy under five-fold cross validation, and then only neurons with a non-zero weight were analysed further.

The mean difference vector was calculated as:

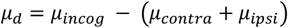

With the corresponding covariance matrix:

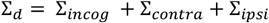

The squared Mahalanobis distance was then calculated as

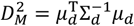

### SVM decoder analysis

In SVM decoder analysis linear SVM decoders were built using the Scikit learn Python module. To decode the current dominant stimulus representation during 30 second congruent or incongruent stimulation, we trained a decoder using responses of all recorded V1 neurons in each experiment to discriminate between responses to a grating of orientation θ presented to the ipsilateral eye, and θ + 90 presented to the contralateral eye. Time averaged responses (from 0 to 2 seconds) to 2 seconds of monocular stimulation were used for model training. Expected performance of the decoder was validated in each animal using hold one out cross validation (Figure 2A), while the model used to assess dominant representation during longer stimulation (i.e. 30 seconds), used all (i.e. 20 in total) short stimulation trials. The decoding model was therefore trained on response data that was independent of that which it was used to classify. For 30 second stimulation trials, responses of neurons were averaged over 1 second bins, and classification was performed on each bin independently (Figure 2B). The fraction of these bins that were classified as either the ipsilaterally or contralaterally presented stimulus was then computed (Figure 2C), and the change in this value was calculated as a function of contralateral stimulus contrast (Figure 2D). To gain a measure of classification confidence we transformed SVM decision values into calibrated probabilities using Platt scaling (*49*). Feature weights learned during SVM model training were multiplied by the neural activity matrix to produce activity vectors of the subpopulation of neurons informative of the presence of the contralaterally or ipsilaterally presented stimulus (Figure 2E). Transitions in classification were defined as a change from one 1-second time bin to the next in the polarity of the SVM decision value (Figure 2E), and average value of the subpopulation was taken by averaging over the −0.5 to 0 seconds before SVM decision switch and the 0 to 1 seconds after (Figure 2G). In order to decoder the dominant movie representation during natural movie stimuli, SVM classifiers were trained to classify 3 second binds of neural activity as having arisen through stimulation with movie_Δ0s_ (presented to ipsilateral eye, see stimulus details above) or movie_Δ10s_ (presented to the contralateral eye). For each pair of 30 second movie, 10 classifiers were therefore trained (Figure 3B). These classifiers were then used to decode the dominant representation when the 2 movies were presented simultaneously (i.e. one to each eye; Figure 3C and D). Analysis of RSC→V1 bouton SVM analysis was performed as per V1 somas with the exception of analysis in which stimulated eye rather than stimulus orientation was used for as a class label.

### Analysis of pupil data

Having extracted pupil diameter from video data (see methods above) calculated stimulus onset aligned pupil diameter traces were then calculated which were baseline subtracted at stimulus onset (Figure 4C). Stimulus onset response amplitude (Figure 4E) was calculated by subtracting pre-stimulus pupil diameter (average of period from −0.5 to 0 prior to stimulus onset) from post-stimulus pupil diameter (average of period from 0 to 2 seconds after stimulus onset). In order to determine if SVM decoder confidence varies as a function of pupil diameter we compared SVM decoder confidence (i.e. Platt scaled decision values) between trials in which the pupil diameter was in the upper or lower 40^th^ percentile of its range (Figure 4F and G).

### Other statistical methods

Statistical analyses were performed using Python (SciPy library), and group average values are presented throughout as mean ± standard error of the mean unless otherwise noted. The statistical significance of comparisons between groups was determined using a two-sided t test or ANOVA unless otherwise noted, and p values <0.05 were considered significant. Precise group sizes were not decided in advance, but approximate group sizes were based on typical sizes used in this field in similar experiments.

| **Analysis** | **Number of mice used** | **Number of neurons or boutons analysed** |
| --- | --- | --- |
| Characterization of binocular incongruent input of V1 somas | 6 mice (7 experiments) | 2176 |
| SVM training and decoding (gratings) | 5 mice (6 experiments) | 1977 |
| SVM training and decoding (natural stimulus) | 6 mice (6 experiments) | 845 |
| SVM training and decoding from RSC→V1 boutons | 4 mice (5 experiments) | 615 |

## Acknowledgements

This work was supported by grants PDI 2019-109285GA-I00, RYC2021-032313-I, PID2022-139822NB-I00, PCI2022-134995-2 to A.R and PRE2020-093128 to M.T. from the Spanish Secretary of Research, Development and Innovation (MINECO), and has received funding from the European Union’s Horizon ERC Grants research and innovation programme under ERC Consolidator grant agreement No. 101088598 (HEU-101088598-DREAMNET) to A.R.

